# orthoSynAssign: refine orthogroups using synteny information

**DOI:** 10.64898/2026.08.10.744007

**Authors:** Cheng-Hung Tsai, Carolina G. Piña Páez, Jason E. Stajich

## Abstract

Accurately identifying orthogroups is crucial for precise phylogenetic reconstruction, but clustering-based methods often generate complex, many-to-many orthogroups that include confounding paralogs. Incorporating synteny offers a robust strategy to refine these clusters into high-granularity, single-copy orthologs. We introduce orthoSynAssign, a user-friendly, high-performance rewrite of the orthogroup refinement tool OrthoRefine, combining an intuitive Python interface with a core computing engine written in Rust. This hybrid architecture ensures straightforward installation, seamless data parsing, and exceptional computational efficiency. Evaluated against the Yeast Gene Order Browser (YGOB) dataset, orthoSynAssign demonstrated outstanding performance, substantially elevating the Area Under the Precision-Recall Curve. Furthermore, multi-threading benchmarks across 193 Eurotiomycetes genomes confirmed strong scalability, drastically reducing execution runtime while maintaining a strictly bounded, thread-independent memory footprint. Ultimately, orthoSynAssign provides a reliable and scalable framework for high-throughput phylogenomic workflows.

## Introduction

Accurately identifying orthogroups is crucial in modern comparative genomics and phylogenetics (Kristensen et al., 2011). These sets of genes, derived from a single ancestral gene via speciation, provide the essential signal for inferring evolutionary history (Sarton-Lohéac et al., 2025). In contrast, paralogs, which arise from gene duplication events, can confound standard phylogenetic reconstruction if they cause gene trees to reflect duplication history rather than true species divergence (Mallatt & Winchell, 2002; Salichos & Rokas, 2013). However, recent research indicates that the inclusion of paralogs can improve phylogenetic inference if modeled correctly (Hellmuth et al., 2015; Smith & Hahn, 2021). Rather than treating duplications as noise to be excluded, the primary challenge is disentangling the orthology of paralogs, a task that requires high-resolution frameworks capable of tracking fine-grained genomic contexts.

Currently, OrthoFinder (Emms & Kelly, 2019) is one of the most widely used tools for de novo orthogroup identification, valued for its high sensitivity and scalability across massive proteomic datasets. However, similarity-based clustering methods like OrthoFinder frequently produce over-aggregated, many-to-many orthogroups containing unresolved paralogs (Tice et al., 2021; Walden & Schranz, 2023). While these broad clusters effectively capture broad evolutionary relationships, they often lack the fine-scale resolution needed for accurate downstream analyses. Resolving these complex, over-aggregated gene families is essential for untangling evolutionary dynamics; without fine-scale orthogroup resolution, over-clustered families introduce noise into single-copy phylogenomics, obscure lineage-specific gene content variation, misrepresent gene duplication and loss events, and distort pangenome architecture (Manzano-Morales et al., 2023; Sarton-Lohéac et al., 2025; Smith et al., 2022).

To resolve these complex clusters, incorporating syntenic conservation—the preservation of relative gene order across chromosomes—provides a robust framework to recover true orthogroups (Catchen et al., 2009; Georgescu et al., 2018; Lovell et al., 2022; Sarkar et al., 2011). Since genomic rearrangements are non-random, the physical proximity of genes provides a robust line of evidence of orthology that sequence similarity alone may fail to capture (Jun et al., 2009; Notebaart et al., 2005). The tool OrthoRefine (Ludwig & Mrázek, 2024) was developed to employ the local syntenic context to refine orthogroups, ensuring that assigned orthologs share a conserved genomic neighborhood and thus mitigating the noise associated with many-to-many clusters.

While OrthoRefine’s elegant design significantly improves the precision of ortholog calls, the original implementation presents substantial usability challenges. As a C++ application, the software requires manual compilation, which is frequently hindered by inconsistencies in compilers and library dependencies across different operating systems. Therefore, successful installation required specialized technical expertise. Beyond installation hurdles, the tool’s data integration framework is notably fragile. In practice, the internal logic frequently fails to correctly map sample information between OrthoFinder-generated orthogroups and their corresponding genomic annotations. These mapping discrepancies lead to recurring execution failures, ultimately compromising the tool’s reliability within automated, high-throughput pipelines.

To address these limitations, orthoSynAssign provides a modern Python and Rust hybrid re-implementation that balances accessibility with high-performance computing. The Python interface offers an intuitive environment for data preprocessing, allowing users to seamlessly interact with and manipulate their dataset objects. Meanwhile, the Rust-implemented core synteny calculation engine maximizes efficiency through rapid computing and a constrained memory footprint, which is essential for large-scale datasets. Ultimately, orthoSynAssign provides a user-friendly installation process and delivers a high-performance framework suitable for high-throughput phylogenomic workflows.

## Materials and methods

### orthoSynAssign workflow

The main interface of orthoSynAssign was developed in Python version 3.9 and above. Prior to analysis, users must convert genome annotations into a sorted, four-column BED format. It is critical that the file basenames (excluding the .bed suffix) precisely match the sample headers in the OrthoFinder-generated orthogroups file. Because synteny analysis relies on genomic coordinates rather than protein sequences used by OrthoFinder for orthogroup detection, the name in the 4th column of these BED files must correspond exactly to the protein IDs represented in the orthogroup file. For genes with multiple isoforms, users may either select a single representative isoform or collapse all isoforms into a single genomic entry; in the latter cases, the fourth column of the BED file should contain a semicolon-delimited concatenation of all associated isoform IDs. For convenience, we provide a utility script to help the GFF-to-BED conversion with multi-isoform concatenation (gff2bed.py). Conceptually, orthoSynAssign operates similarly to OrthoRefine, using a local-synteny-driven workflow summarized in Fig. 1. Within the Python refinement module, each gene entry is instantiated as a Gene object and cross-referenced with its corresponding Genome and Orthogroup objects. For genes with multiple isoforms (n > 1), the module creates n+1 objects: one for each individual isoform and a single representative object for the gene as a whole. Only these representative objects are used for the synteny calculations. Following refinement, results are mapped back to the original isoform objects to accurately reflect protein-level orthology. This object-oriented architecture provides a flexible environment for users to explore, visualize, and customize their analyses within interactive platforms such as Jupyter Notebooks, Google Colab, or PyCharm.

**Figure 1.**
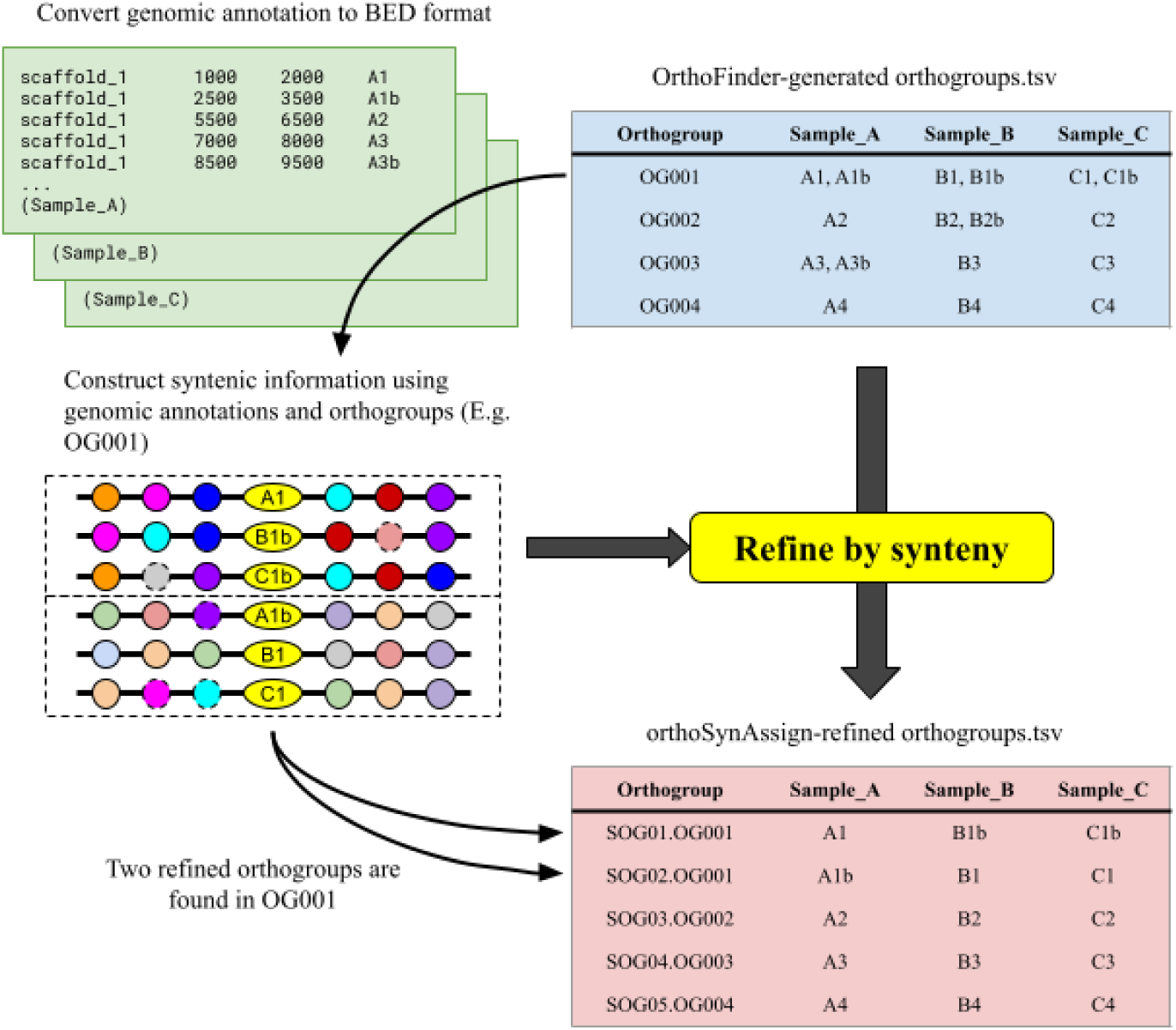
Flowchart of orthoSynAssign. Genomic annotations for multiple samples (e.g. Sample_A, Sample_B, Sample_C) are converted into BED format and cross-referenced with sequence-similarity orthogroups generated by OrthoFinder (OrthoFinder-generated orthogroups.tsv). By mapping these orthogroups back onto their syntenic context, orthoSynAssign constructs syntenic blocks. The pipeline then evaluates gene arrangements to resolve ambiguous or multi-copy assignments within a single sequence-similarity cluster (e.g., separating OG001 into distinct syntenic groups). The final output (orthoSynAssign-refined orthogroups.tsv) yields refined syntenic orthogroups (SOGs) that reflect true positional orthology across genomes.

While Python offers significant flexibility, its memory management can be a bottleneck for high-throughput, large-scale synteny analysis. To optimize the workflow and minimize the memory footprint, we implemented the core synteny calculation engine in Rust (2021 Edition), a programming language which is well known for its memory safety and C/C++ comparable execution speed (Matsakis & Klock, 2014) and interacts with Python interface using PyO3 crate (Johnson & Hodson, 2025). More importantly, this implementation bypasses Python’s Global Interpreter Lock (GIL), unlocking the full potential of multi-core systems for true parallel processing. Ultimately, the syntenically refined orthogroups are exported in the same tsv format as the initial OrthoFinder input to ensure downstream compatibility.

To resolve orthogroup ambiguity and refine gene assignments across multiple genomes, orthoSynAssign introduces two key algorithmic advances over OrthoRefine: pre-analysis tandem repeat collapsing and graph-based Disjoint Set Union (DSU) clustering. First, whereas OrthoRefine handles local gene duplications heuristically by identifying adjacent HOG members during pairwise window scans (together1/together2 vectors) and concatenating their locus tags into comma-separated strings without collapsing them, orthoSynAssign explicitly collapses tandem duplicate clusters into single syntenic units prior to sliding-window calculation, preventing local window span inflation and uninformative orthogroup matching. Second, following the pairwise synteny evaluation, OrthoRefine performs multi-species clustering through repeated linear searches over dynamic, multi-dimensional array structures; in contrast, orthoSynAssign models the pairwise syntenic gene pairs as edges in an undirected graph and executes DSU clustering (Cormen & Leiserson, 2022) with two-pass path compression in near-constant amortized time *O*(α(*N*)) (Tarjan, 1975), mapping connected components back to genome-specific locus identifiers in a single post-processing pass to preserve genuine orthology while minimizing memory overhead.

To complement the refinement process, we developed a companion script for result visualization (orthosynassign-vis). This script utilizes the same BED annotation files, the orthogroups files (pre- and post-refinement), and the specific ID(s) of the orthogroups of interest. It generates a figure that delineates the syntenic relationships within a window, illustrating the genomic context of all genes within the refined orthogroup.

### Dataset for benchmark

Two datasets were used to benchmark orthoSynAssign. The first comprises 20 yeast species, as utilized in Yeast Gene Order Browser (YGOB) Version 7 (Byrne & Wolfe, 2005). In addition to the genomes and annotations, YGOB also provides rigorously verified orthogroups (“pillars”) spanning both pre- and post-whole genome duplication (WGD) events associated with the divergence of *Saccharomyces* species and some closely related species that do not undergo WGD. These well-curated evolutionary relationships serve as an ideal gold-standard resource for validating synteny-based orthology assignment.

To evaluate computational efficiency and scalability, we benchmarked the tool using *Eurotiomycetes* (TaxID: 147545), a major class of *ascomycete* fungi represented by a vast and diverse collection of genomic data. Using the NCBI Datasets command-line tool (O’Leary et al., 2024), we retrieved all annotated, high-quality reference assemblies filtered for chromosome-, complete-, or scaffold-level resolution:

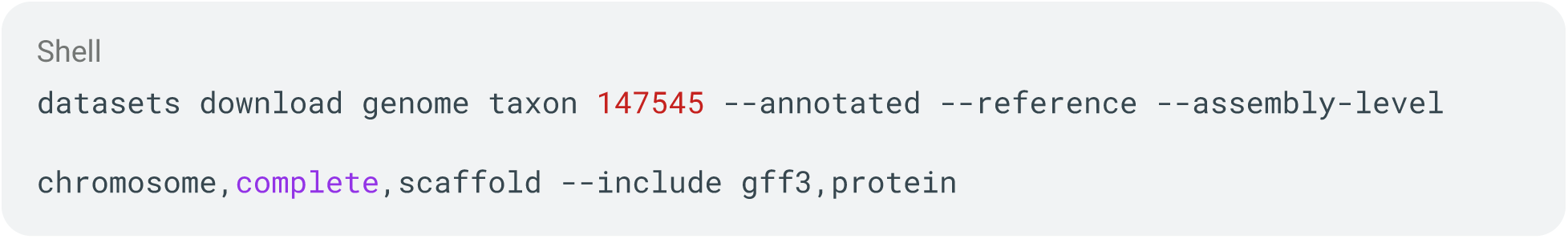

This query yielded a target dataset of 193 genomes, complete with structural annotations GFF and protein sequences. Because orthoSynAssign relies fundamentally on exhaustive pairwise comparisons, an input dataset of this scale introduces a steep combinatorial expansion. Processing these 193 genomes poses a substantial computational challenge, serving as an ideal stress test for the tool’s peak memory consumption, runtime scalability, and overall algorithmic efficiency. For all benchmark evaluations across both datasets, orthoSynAssign was executed using its default settings: a synteny ratio threshold of ≥ 0.5 within an 8-gene window.

### Performance evaluation

The YGOB orthogroups (“pillars”) was utilized as the ground truth to evaluate the performance of orthoSynAssign. Because YGOB is a purely synteny-based framework that does not account for sequence similarity, its orthogroups often contain divergent proteins that are excluded during OrthoFinder’s sequence-similarity screening. To ensure a fair comparison, we first ran OrthoFinder v2.5.5 with “-S diamond” and “-og” flags to establish the sequence-based baseline. We then filtered the YGOB orthogroups to retain only the proteins present in these OrthoFinder results, eliminating divergent proteins that would otherwise artificially inflate the false-negative rate. These filtered orthogroups served as the standardized ground truth to benchmark both the raw OrthoFinder orthogroups and those refined by orthoSynAssign. The performance evaluation was conducted at two scales: the overall-orthogroup level and the within-orthogroup level.

The overall-orthogroup level assessment evaluates how many complete orthogroups are correctly identified relative to the ground truth (Fig. S1). First, the Jaccard similarity is calculated between all possible orthogroup pairs from the testing method and the ground truth based on their shared protein compositions. This generates a m × n Jaccard similarity matrix, where m is the number of YGOB orthogroups and n is the number of orthogroups from the testing method. We then employ a strict classification strategy where a pair is designated as a true positive (TP) if and only if it (1) established as a reciprocal best match within the matrix, and (2) its Jaccard similarity score meets or exceeds the predefined cutoff. Consequently, ground-truth orthogroups that fail to establish a reciprocal best match or fall below the Jaccard cutoff are classified as false negatives (FN). Conversely, orthogroups from the testing method that either lack a reciprocal counterpart or fail to pass the Jaccard threshold are classified as false positives (FP). By iteratively adjusting this Jaccard cutoff, we can generate a precision-recall curve and comprehensively evaluate performance by calculating the area under the curve (AUC).

The within-orthogroup level assessment examines the precision of individual protein assignments within these matched orthogroup pairs. We construct a distribution histogram for each method using the maximum Jaccard scores from the reciprocal best match step, prior to applying any cutoff filtering. This allows us to visualize and compare how closely the internal protein compositions of the predicted orthogroups align with their respective ground-truth counterparts.

To evaluate how orthogroup refinement impacts species tree inference, we compared 2,406 raw single-copy orthogroups identified from YGOB dataset prior to refinement and 1,162 strict single-copy orthogroups derived after orthoSynAssign refinement. For each dataset, sequence alignments were generated using MAFFT v7.505 (Katoh & Standley, 2013), and individual gene trees were inferred using FastTree v2.1.11 (Price et al., 2010). Species trees were subsequently reconstructed using ASTRAL-Pro 3 v1.19.3.6 (Zhang et al., 2025). To evaluate node confidence across the species trees, we calculated two complementary support metrics: gene concordance factors (gCF) using IQ-TREE v2.4.0 (Minh et al., 2020), which quantify the proportion of individual gene trees that support each clade, and local quartet support values generated by ASTRAL-Pro 3.

### Computational Efficiency Benchmarking

The Eurotiomycetes dataset was utilized to evaluate the computational efficiency of orthoSynAssign. To generate the initial orthogroups, OrthoFinder was executed with the same settings described above. The computational efficiency of orthoSynAssign was benchmarked across varying CPU allocations (configured via the “--threads/-t” parameters) and resource consumption was monitored using the Python package memory-profiler v0.61.0 (Pedregosa, 2022). To ensure consistency and reproducibility, each thread configuration was benchmarked three times under identical conditions on a dedicated high-performance computing cluster node equipped with an AMD EPYC 9634 84-Core Processor (2.25 GHz base clock, up to 3.7 GHz boost clock).

### Data visualization

Data visualization was primarily performed using the Python package matplotlib (Hunter, 2007) and its high-level interface seaborn (Waskom, 2021). Comparative phylogenetic trees were visualized using the R package phytools 2.5.2 (Revell, 2024).

## Results and discussion

### orthoSynAssign successfully splitting over-aggregating orthogroups using synteny information

One of the most direct methods to assess the refinement efficacy of orthoSynAssign is to evaluate the average protein count per sample across all orthogroups. An average count of exactly one protein per sample indicates a strict single-copy ortholog, the hallmark of true 1:1 orthology. Accordingly, we calculated this metric for the raw OrthoFinder orthogroups identified from the YGOB dataset and compared it against the orthoSynAssign-refined result. Prior to refinement, a substantial number of orthogroups exhibited an elevated average protein count per sample. Following orthoSynAssign refinement, however, every orthogroup contained fewer than three proteins per sample, with the distribution’s peak shifting decisively into the 0–1 interval (Fig. 2A). This result was consistent with the Eurotiomycetes dataset (Fig. S2). This dramatic contraction demonstrates that orthoSynAssign successfully resolves chimeric clusters by splitting mixed orthogroups and segregating paralogous sequences.

**Figure 2.**
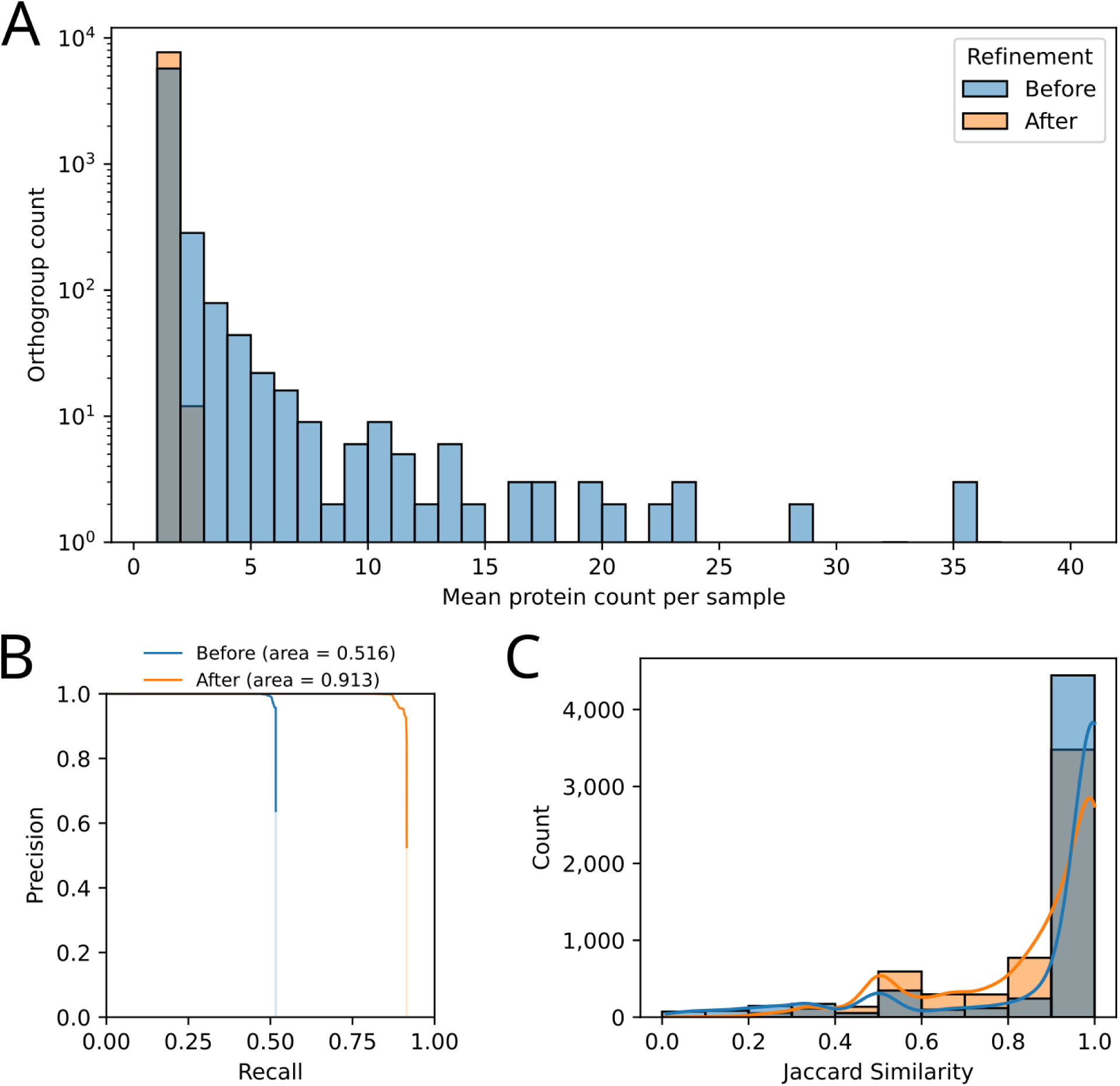
Orthogroup classification performance metrics before and after orthoSynAssign refinement benchmarked with YGOB dataset. (A) Histogram showing the distribution of orthogroup counts relative to the mean protein count per sample. Refinement (orange) significantly condenses inflated orthogroup sizes toward single-copy or low-copy expectations compared to pre-refined baseline (blue). (B) Precision-Recall curve highlighting performance optimization. The area under the curve increases from 0.516 (before refinement, blue) to 0.913 (after refinement, orange). (C) Distribution of Jaccard similarity relative to the ground truth baseline. Both prior to refinement (blue) and following orthoSynAssign refinement (orange), the distributions exhibit a peak at a Jaccard score of 1.0. This score represents an identical gene-for-gene composition, indicating precise alignment with the ground-truth baseline.

To comprehensively evaluate the performance of orthoSynAssign, we utilized the YGOB-defined “pillars” (orthogroups) as the ground truth to calculate classification metrics. At the overall-orthogroup level, we computed the Area Under the Precision-Recall Curve (AUPRC) for both the raw OrthoFinder orthogroups and the orthoSynAssign-refined results. Following refinement, the AUPRC improved significantly from 0.516 to 0.913 (Fig. 2B). This dramatic increase demonstrates that the refined orthogroups are highly congruent with the curated YGOB reference, largely driven by orthoSynAssign successfully dismantling over-aggregated OrthoFinder orthogroups and segregating confounding paralogs.

At the within-orthogroup level, we evaluated internal consistency by analyzing the Jaccard similarity scores of reciprocally matched orthogroups before and after refinement against the YGOB reference dataset. In both cases, the primary peak of the Jaccard similarity distribution was centered at 1.0, confirming strong baseline alignment (Fig. 2C). Following refinement, however, the peak showed a slight reduction in height and a minor broadening. This subtle shift points to occasional over-splitting, where true orthologous members were erroneously partitioned into separate orthogroups. Nevertheless, the refined orthogroups retained high compositional accuracy, demonstrating close structural alignment with the YGOB reference. Visual inspection of individual split orthogroups confirmed that orthoSynAssign successfully segregated genes according to the defined threshold, yielding sub-clusters with distinctly higher syntenic purity than the unrefined parent orthogroups (Fig. 3 and S3).

**Figure 3.**
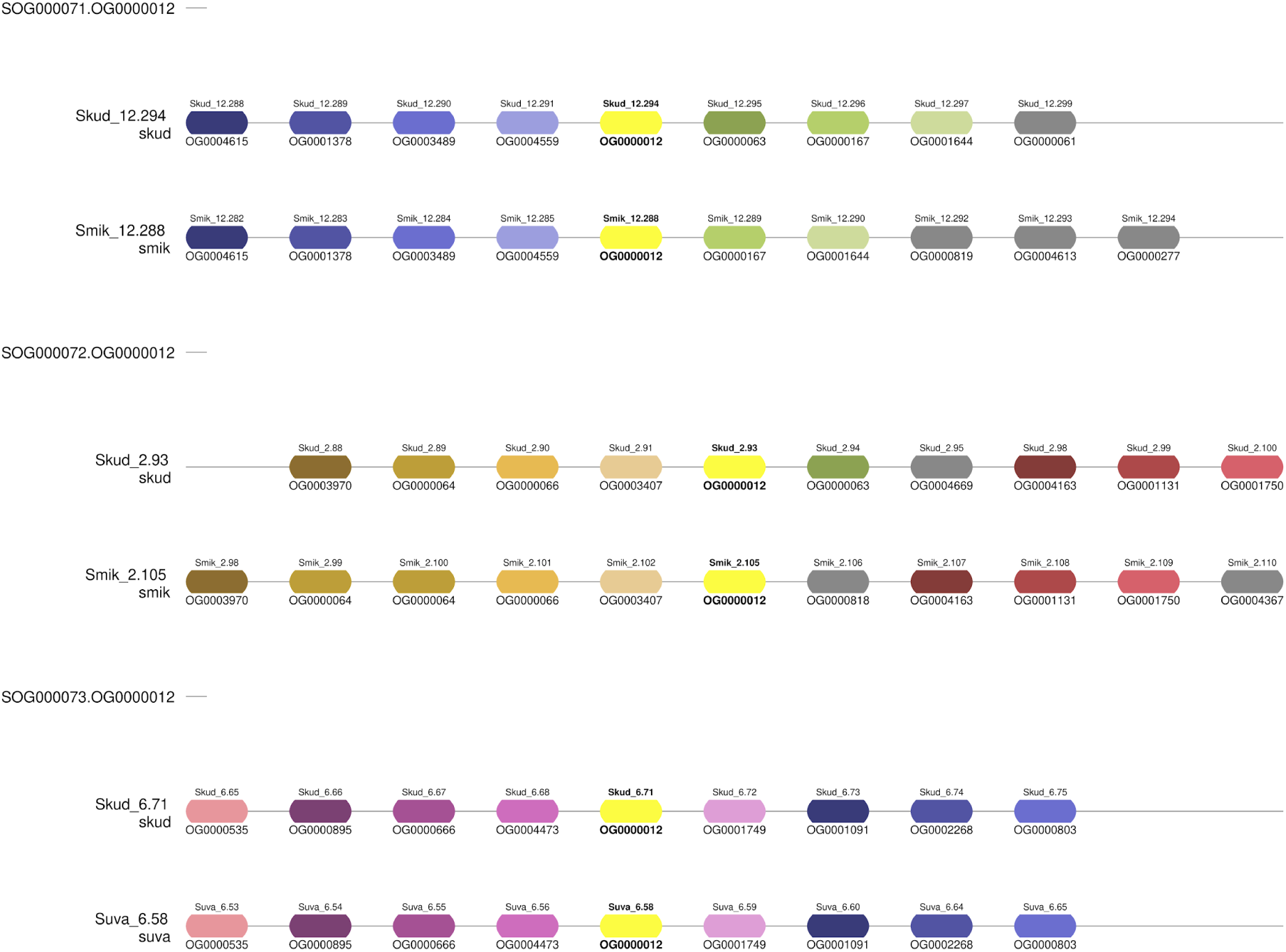
Resolution of paralogous genomic contexts within orthogroup OG0000012. Syntenic alignment and partitioning of raw orthogroup OG0000012 into three distinct sub-orthogroups (SOG000071–SOG000073) by orthoSynAssign. Target genes are shown in bold text and colored in bright yellow, flanked by neighboring genes colored by orthogroup assignment. orthoSynAssign successfully separated loci based on local genomic context (synteny ratio ≥ 0.5 within an 8-gene window).

To assess the practical impact of synteny-guided orthogroup refinement on phylogenetic inference, we evaluated species tree reconstruction using the YGOB dataset by comparing species trees generated from single-copy orthogroups before and after orthoSynAssign refinement. Both datasets yielded identical tree topologies across all YGOB yeast species (Fig. S4), consistently recovering the established Post-WGD, ZT, and KLE clades (Wolfe, 2015). However, the refined dataset demonstrated a systematic improvement in phylogenetic support metrics across internal nodes. Local quartet support values from ASTRAL-Pro 3 consistently shifted upward, accompanied by modest increases in gene concordance factors (gCF). Although raw single-copy identification captures a larger volume of candidate orthogroups, the lower support values in the unrefined tree suggest the presence of residual paralogy or locus-specific noise. By filtering out these confounding signals, orthoSynAssign enhances gene tree concordance and sharpens phylogenetic signals without distorting underlying evolutionary relationships.

### orthoSynAssign demonstrates high computational efficiency and a minimized memory footprint

We benchmarked the computational efficiency and scalability of orthoSynAssign by refining OrthoFinder orthogroups across a dataset of 193 Eurotiomycetes genomes evaluated on 1, 2, 4, 8, 16, and 32 CPU cores. Overall, the computational runtime of orthoSynAssign demonstrates strong multi-threading scalability (Fig. 4). Using a single CPU core, the total execution time was approximately 14 minutes. Increasing the core allocation yielded near-linear speedups up to 8 CPUs, dropping the runtime sharply to roughly 3 minutes. Beyond 16 cores, the runtime curve begins to plateau, approaching a floor of approximately 1.5 minutes at 32 CPUs (Table S1).

**Figure 4.**
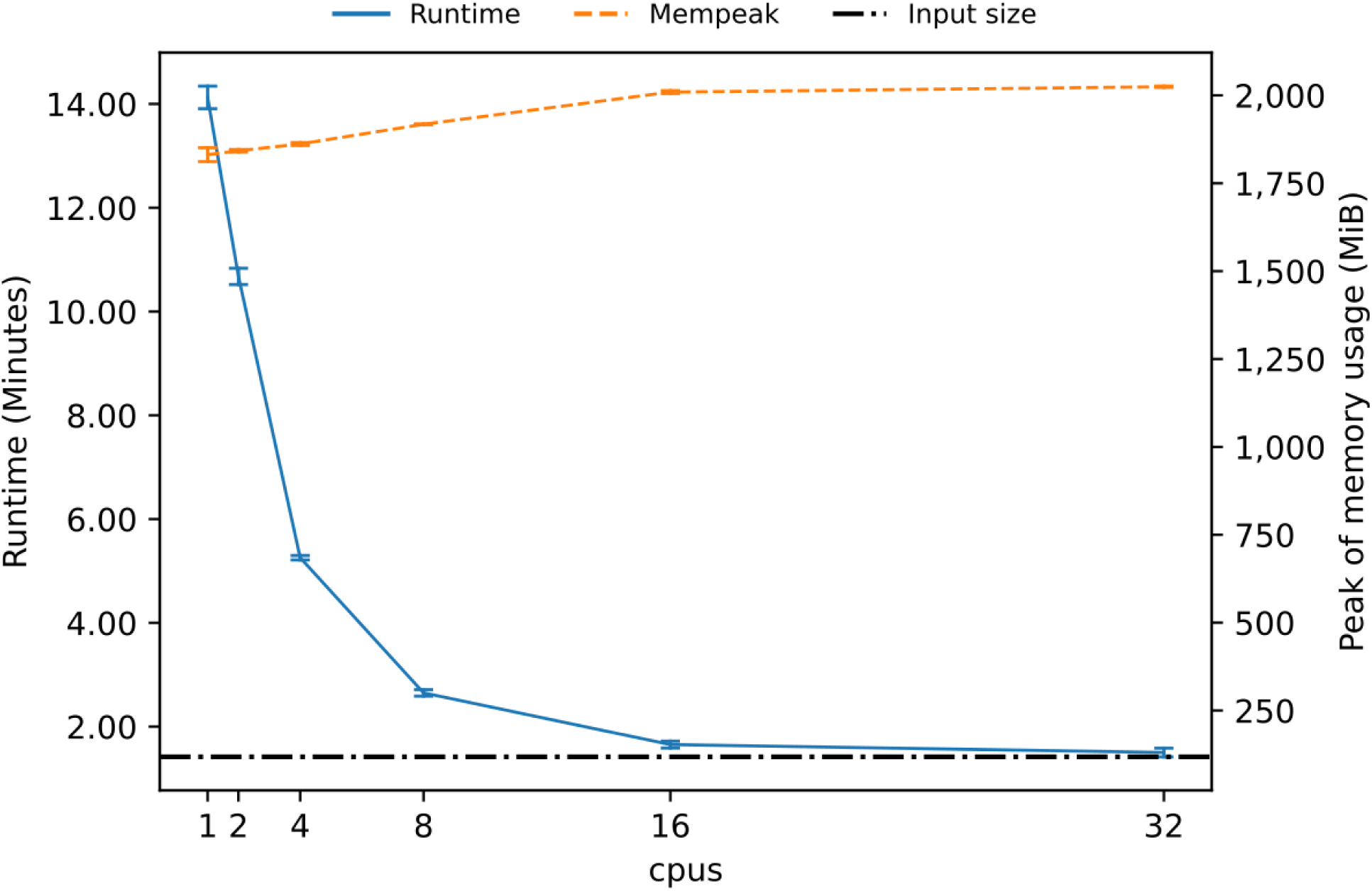
Computational benchmarking of orthoSynAssign across varying CPU allocations. Execution runtime in minutes (solid blue line, left y-axis) and peak memory usage in MiB (dashed orange line, right-axis) are plotted against allocated CPU threads (1 to 32). The horizontal black dash-dotted line marks the input file size. The pipeline scales efficiently, reducing runtime from 14 minutes to 1.5 minutes at 16 or more CPUs, while keeping memory usage stable at approximately 2,000 MiB. Error bars represent standard deviation across three replicates.

In terms of memory consumption, orthoSynAssign exhibits an exceptionally stable and efficient profile across all CPU configurations. On a single CPU core, peak memory usage is approximately 1,800 MiB. As the number of threads increases from 1 to 32, peak memory usage rises only marginally, stabilizing at an upper limit of roughly 2,000 MiB. This indicates that the total memory overhead remains strictly constrained and predictable regardless of core allocation. This minimal memory expansion demonstrates that the Rust-based calculation engine avoids significant parallel data duplication, maintaining high memory safety and efficiency even under maximal multi-threaded scaling.

## Supporting information

Supplementary File 1. Metadata, BED files and Orthogroup tables for the YGOB dataset generated using OrthoFinder, YGOB pillars and orthoSynAssign

Supplementary File 2. BED files and Orthogroup tables for the Eurotiomycetes dataset generated using OrthoFinder and orthoSynAssign

## Data availability

The orthoSynAssign package is available at https://github.com/stajichlab/orthoSynAssign, https://anaconda.org/bioconda/orthosynassign and Zenodo doi: 10.5281/zenodo.18762979. The analyses described in this study were performed using the release v1.3.1 archived as doi: 10.5281/zenodo.21844775. The GFF-to-BED conversion script is available at https://github.com/stajichlab/orthoSynAssign/blob/main/misc/gff2bed.py. The BED files, orthogroup tables for YGOB and Eurotiomycetes datasets are available at Supplementary File 1 and 2.

## Author contributions

C.-H.T.: Conceptualization, Methodology, Software, Validation, Formal analysis, Data Curation, Writing – original draft, Visualization. C.P.P.: Conceptualization, Data Curation, Writing – review & editing. J.E.S.: Conceptualization, Software, Methodology, Resources, Supervision, Project administration, Funding acquisition, Writing – review & editing. All authors read and approved the final manuscript.

## Acknowledgements

This work was supported by National Science Foundation (NSF) grants DEB-1441715 and EF-2125066 (to J.E.S., C.P.P., and C.-H.T.), subawards under National Institutes of Health (NIH) grants R01AI130128 and R01AI127548 (to J.E.S., C.P.P., and C.-H.T.), and U.S. Department of Agriculture, National Institute of Food and Agriculture (USDA-NIFA) grants 2020-70029-33202 and CA-R-PPA-211-5062-H (to C.-H.T. and J.E.S.). J.E.S. is a CIFAR Fellow in the program Fungal Kingdom: Threats and Opportunities. Computational analyses and data storage were supported by the High-Performance Computing Cluster (HPCC) at UC Riverside, funded by NSF grants (MRI-2215705, MRI-1429826) and NIH grant 1S10OD016290-01A1.

