## Supplementary Tables and Figures for "orthoSynAssign: refine orthogroups using synteny information"

Supplementary Table 1. Computational benchmarking of orthoSynAssign across varying CPU allocations.

| CPUs | Runtime (min) | | Peak of mem usage (MiB) | |
|---|---|---|---|---|
| | Mean | Standard deviation | Mean | Standard deviation |
| 1 | 14.13 | 0.22 | 1,831.35 | 19.57 |
| 2 | 10.68 | 0.16 | 1,842.40 | 3.49 |
| 4 | 5.26 | 0.04 | 1,861.56 | 3.97 |
| 8 | 2.65 | 0.06 | 1,917.66 | 1.08 |
| 16 | 1.65 | 0.07 | 2,009.16 | 4.02 |
| 32 | 1.50 | 0.08 | 2,024.31 | 1.77 |

Mean, Standard deviation and Standard error of mean are calculated across three trials

### Supplementary Figures

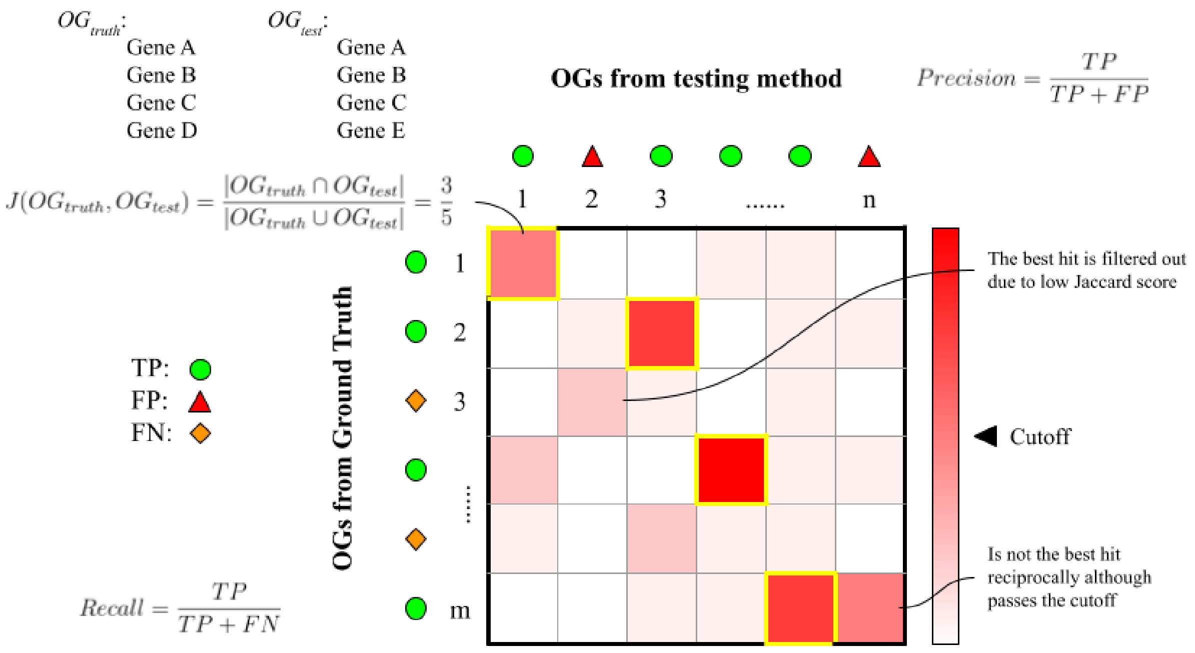


Supplementary Figure 1. Orthogroup classification strategy using a Jaccard similarity matrix. A Jaccard similarity matrix (m × n) is generated by calculating the shared protein composition between orthogroups from the ground truth (YGOB, rows) and a given testing method (columns). A strict classification strategy designates a pair as a true positive (green circles) if it forms a reciprocal best match that meets or exceeds a predefined Jaccard similarity cutoff (black arrowhead). Orthogroups from either the ground truth or the testing method that lack a valid reciprocal counterpart or fall below the threshold are classified as false negatives (orange diamonds) and false positives (red triangles), respectively.

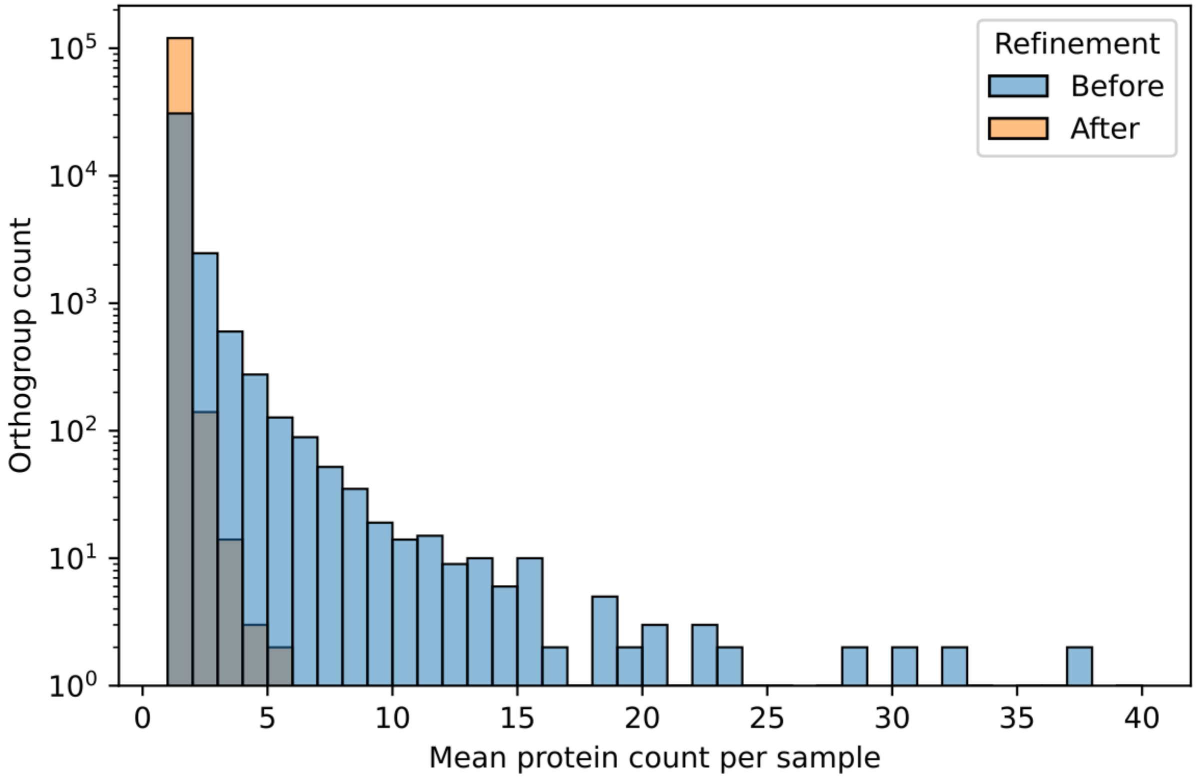


Supplementary Figure 2. Distribution of orthogroup sizes before and after refinement in Eurotiomycetes. Histogram showing orthogroup gene count distributions relative to the mean protein count per sample across Eurotiomycetes. Dataset refinement (orange) significantly reduces inflated orthogroup sizes toward single- or low-copy expectations compared to the pre-refined baseline (blue).

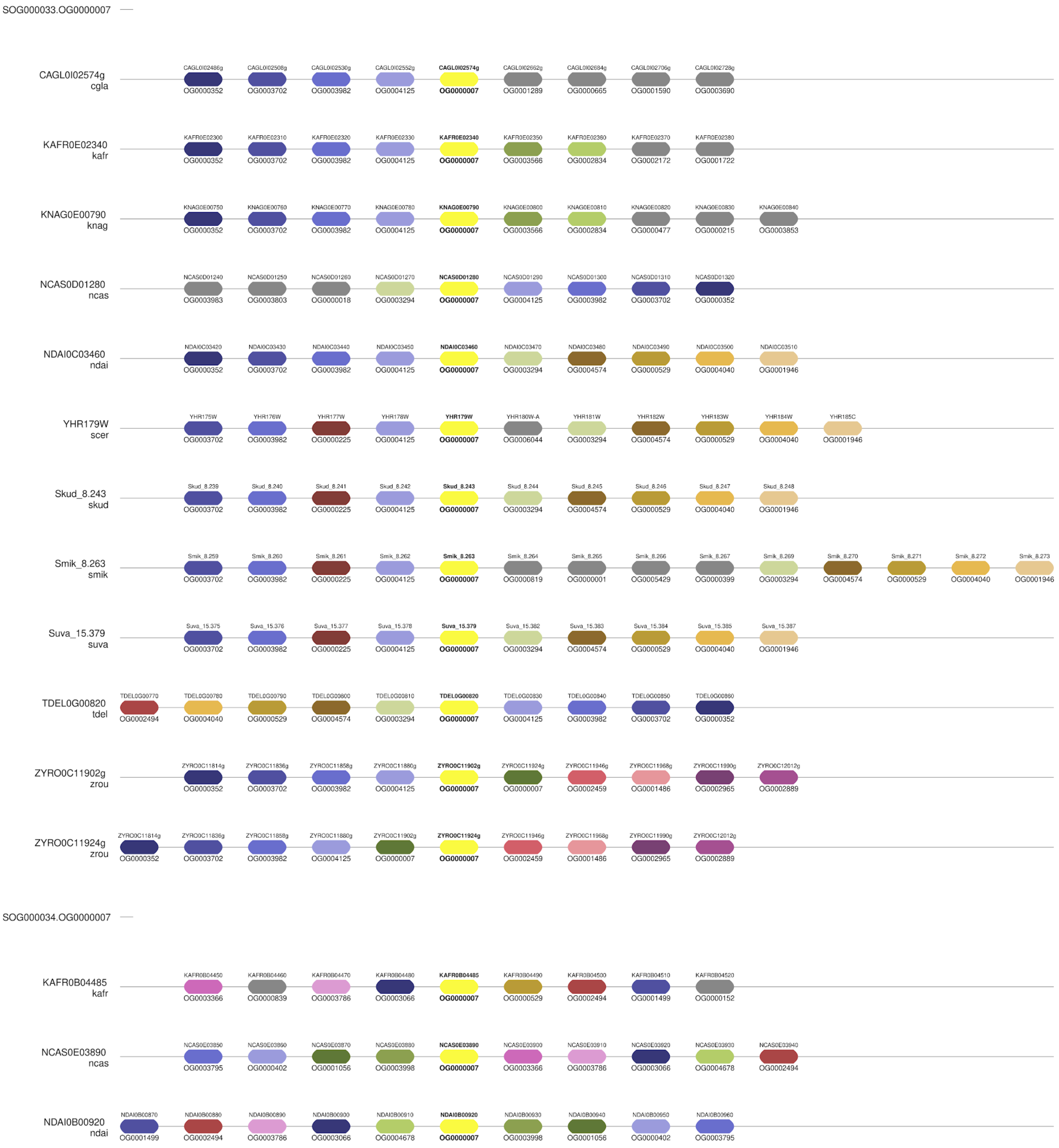

SOG000033.OG0000007
CAGL0I02574g
cgla
CAGL0I02486g CAGL0I02508g CAGL0I02530g CAGL0I02552g CAGL0I02574g CAGL0I02662g CAGL0I02684g CAGL0I02706g CAGL0I02728g
OG0000352 OG0003702 OG0003982 OG0004125 OG0000007 OG0001289 OG0000665 OG0001590 OG0003690
KAFR0E02340
kafr
KAFR0E02300 KAFR0E02310 KAFR0E02320 KAFR0E02330 KAFR0E02340 KAFR0E02350 KAFR0E02360 KAFR0E02370 KAFR0E02380
OG0000352 OG0003702 OG0003982 OG0004125 OG0000007 OG0003566 OG0002834 OG0002172 OG0001722
KNAG0E00790
knag
KNAG0E00750 KNAG0E00760 KNAG0E00770 KNAG0E00780 KNAG0E00790 KNAG0E00800 KNAG0E00810 KNAG0E00820 KNAG0E00830 KNAG0E00840
OG0000352 OG0003702 OG0003982 OG0004125 OG0000007 OG0003566 OG0002834 OG0000477 OG0000215 OG0003853
NCAS0D01280
ncas
NCAS0D01240 NCAS0D01250 NCAS0D01260 NCAS0D01270 NCAS0D01280 NCAS0D01290 NCAS0D01300 NCAS0D01310 NCAS0D01320
OG0003983 OG0003803 OG0000018 OG0003294 OG0000007 OG0004125 OG0003982 OG0003702 OG0000352
NDAI0C03460
ndai
NDAI0C03420 NDAI0C03430 NDAI0C03440 NDAI0C03450 NDAI0C03460 NDAI0C03470 NDAI0C03480 NDAI0C03490 NDAI0C03500 NDAI0C03510
OG0000352 OG0003702 OG0003982 OG0004125 OG0000007 OG0003294 OG0004574 OG0000529 OG0004040 OG0001946
YHR179W
scer
YHR175W YHR176W YHR177W YHR178W YHR179W YHR180W-A YHR181W YHR182W YHR183W YHR184W YHR185C
OG0003702 OG0003982 OG0000225 OG0004125 OG0000007 OG0006044 OG0003294 OG0004574 OG0000529 OG0004040 OG0001946
Skud_8.243
skud
Skud_8.239 Skud_8.240 Skud_8.241 Skud_8.242 Skud_8.243 Skud_8.244 Skud_8.245 Skud_8.246 Skud_8.247 Skud_8.248
OG0003702 OG0003982 OG0000225 OG0004125 OG0000007 OG0003294 OG0004574 OG0000529 OG0004040 OG0001946
Smik_8.263
smik
Smik_8.259 Smik_8.260 Smik_8.261 Smik_8.262 Smik_8.263 Smik_8.264 Smik_8.265 Smik_8.266 Smik_8.267 Smik_8.269 Smik_8.270 Smik_8.271 Smik_8.272 Smik_8.273
OG0003702 OG0003982 OG0000225 OG0004125 OG0000007 OG0000819 OG0000001 OG0005429 OG0000399 OG0003294 OG0004574 OG0000529 OG0004040 OG0001946
Suva_15.379
suva
Suva_15.375 Suva_15.376 Suva_15.377 Suva_15.378 Suva_15.379 Suva_15.382 Suva_15.383 Suva_15.384 Suva_15.385 Suva_15.387
OG0003702 OG0003982 OG0000225 OG0004125 OG0000007 OG0003294 OG0004574 OG0000529 OG0004040 OG0001946
TDEL0G00820
tdel
TDEL0G00770 TDEL0G00780 TDEL0G00790 TDEL0G00800 TDEL0G00810 TDEL0G00820 TDEL0G00830 TDEL0G00840 TDEL0G00850 TDEL0G00860
OG0002494 OG0004040 OG0000529 OG0004574 OG0003294 OG0000007 OG0004125 OG0003982 OG0003702 OG0000352
ZYRO0C11902g
zrou
ZYRO0C11814g ZYRO0C11836g ZYRO0C11858g ZYRO0C11880g ZYRO0C11902g ZYRO0C11924g ZYRO0C11946g ZYRO0C11968g ZYRO0C11990g ZYRO0C12012g
OG0000352 OG0003702 OG0003982 OG0004125 OG0000007 OG0000007 OG0002459 OG0001486 OG0002965 OG0002889
ZYRO0C11924g
zrou
ZYRO0C11814g ZYRO0C11836g ZYRO0C11858g ZYRO0C11880g ZYRO0C11902g ZYRO0C11924g ZYRO0C11946g ZYRO0C11968g ZYRO0C11990g ZYRO0C12012g
OG0000352 OG0003702 OG0003982 OG0004125 OG0000007 OG0000007 OG0002459 OG0001486 OG0002965 OG0002889
SOG000034.OG0000007
KAFR0B04485
kafr
KAFR0B04450 KAFR0B04460 KAFR0B04470 KAFR0B04480 KAFR0B04485 KAFR0B04490 KAFR0B04500 KAFR0B04510 KAFR0B04520
OG0003366 OG0000839 OG0003786 OG0003066 OG0000007 OG0000529 OG0002494 OG0001499 OG0000152
NCAS0E03890
ncas
NCAS0E03850 NCAS0E03860 NCAS0E03870 NCAS0E03880 NCAS0E03890 NCAS0E03900 NCAS0E03910 NCAS0E03920 NCAS0E03930 NCAS0E03940
OG0003795 OG0000402 OG0001056 OG0003998 OG0000007 OG0003366 OG0003786 OG0003066 OG0004678 OG0002494
NDAI0B00920
ndai
NDAI0B00870 NDAI0B00880 NDAI0B00890 NDAI0B00900 NDAI0B00910 NDAI0B00920 NDAI0B00930 NDAI0B00940 NDAI0B00950 NDAI0B00960
OG0001499 OG0002494 OG0003786 OG0003066 OG0004678 OG0000007 OG0003998 OG0001056 OG0000402 OG0003795

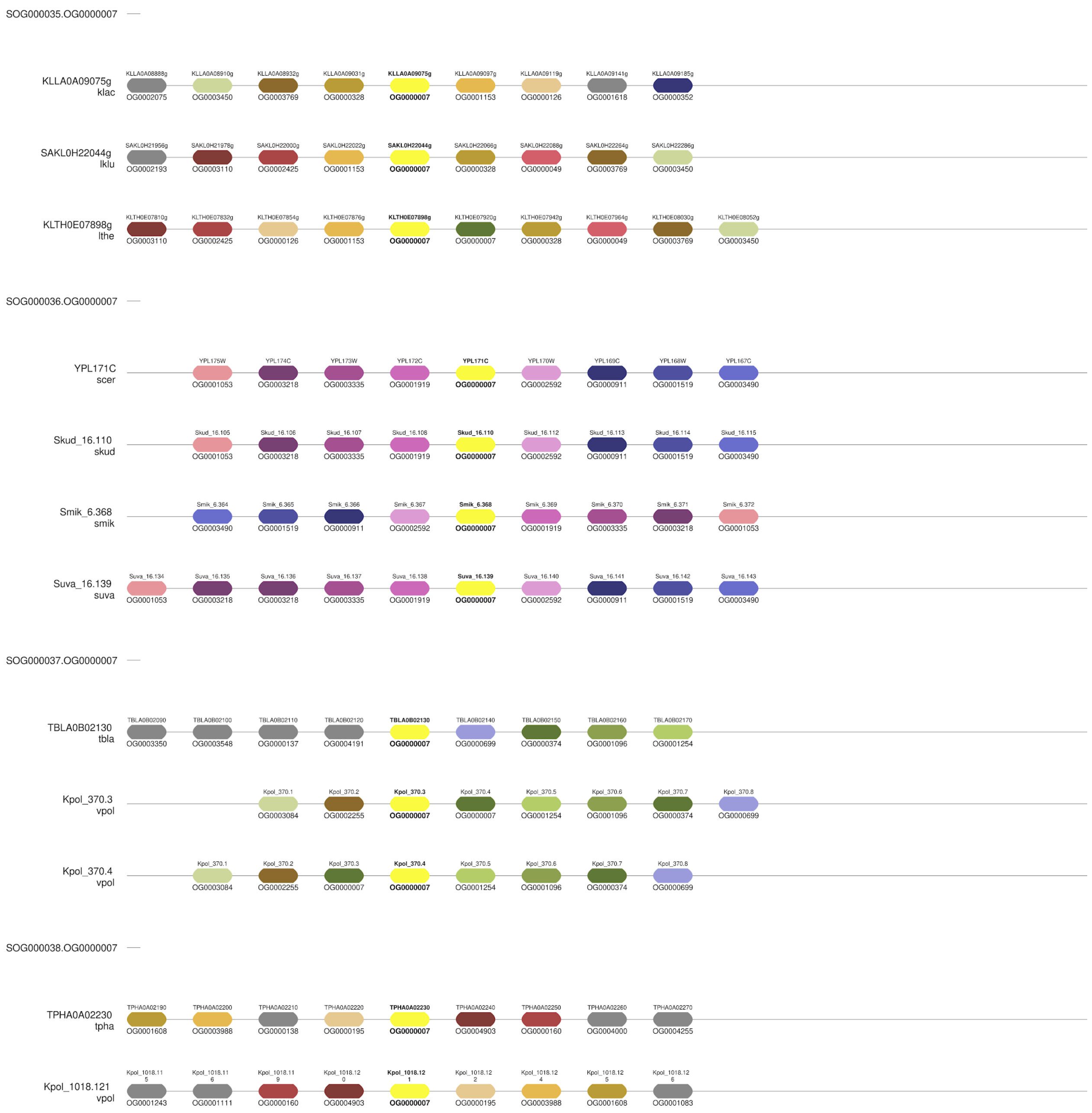


Supplementary Figure 3. Syntenic partitioning of orthogroup OG0000007. Resolution of raw orthogroup OG0000007 into six syntenically pure sub-orthogroups (SOG000033–SOG000038) by orthoSynAssign. Target genes are shown in bold text and colored in bright yellow, flanked by neighboring genes colored by orthogroup assignment. orthoSynAssign successfully separated loci based on local genomic context (synteny ratio $\geq 0.5$ within an 8-gene window).

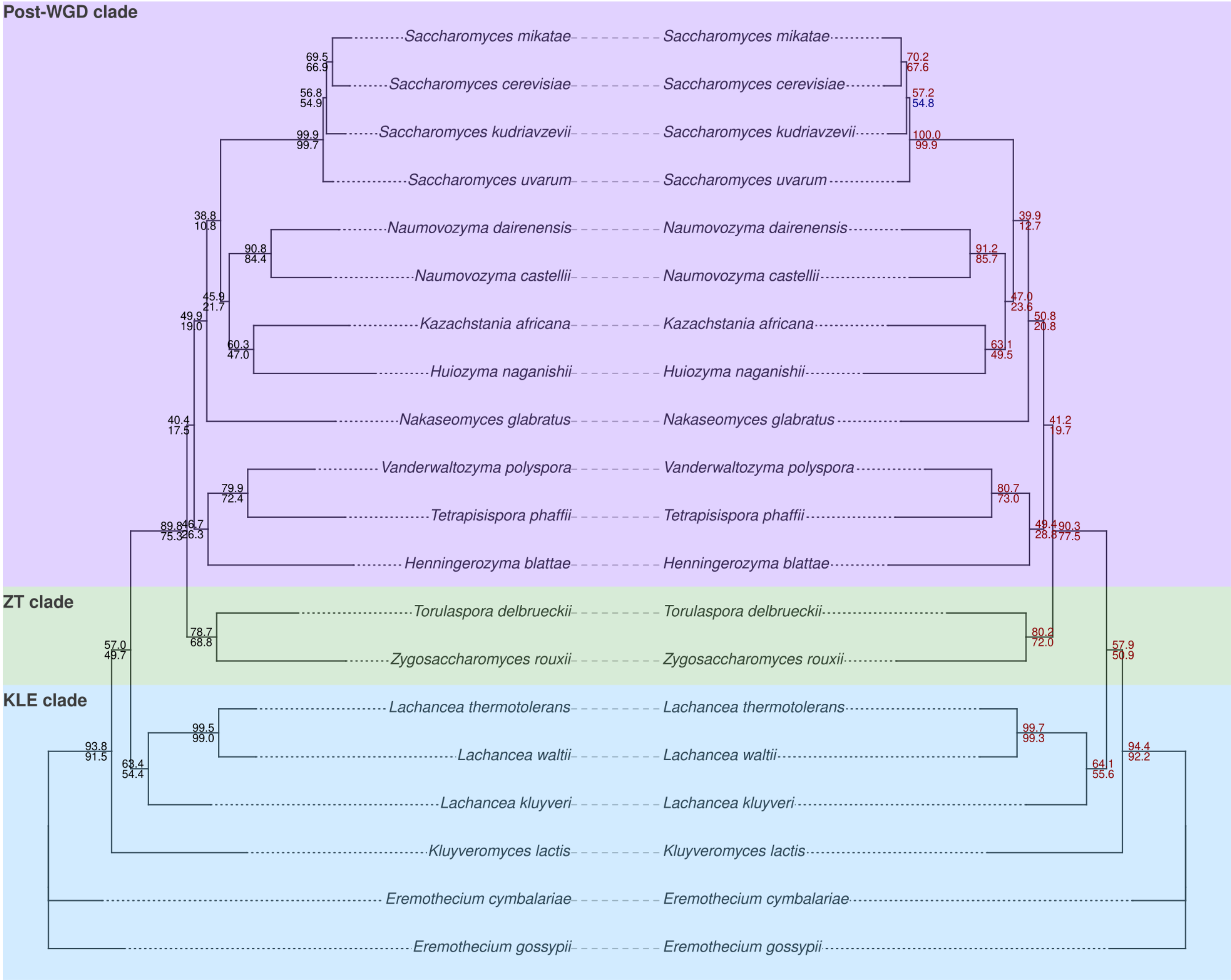


Supplementary Figure 4. Impact of synteny-based orthogroup refinement on species tree topology and node support in yeasts. Tanglegram comparing species trees reconstructed from 20 yeast species in the Yeast Gene Order Browser (YGOB) dataset before (left) and after (right) orthoSynAssign refinement. Shaded background blocks detonate major evolutionary lineages: Post-Whole Genome Duplication (Post-WGD; purple), *Zygosaccharomyces/Torulaspora* (ZT; green), and *Kluyveromyces/Lachancea/Eremothecium* (KLE; blue). Numbers at internal nodes represent local quartet support values from ASTRAL-Pro 3 (top) and gene concordance factors (gCF, bottom) calculated in IQ-TREE, shown in black font on the left tree and color-coded on the right tree to indicate higher (red) or lower (blue) values relative to the unrefined baseline.
